# BRDriver: Breast Cancer Driver Gene Predictor

**DOI:** 10.1101/2023.01.09.523362

**Authors:** Rajitha Perera, Roshan Perera

## Abstract

The breast cancer mortality rate is high in developing countries as the early detection of cancer is deficient in patients. The identification of the genes that drive cancer is one of the major approaches in the early detection of breast cancer. Several computational tools have been developed to predict the cancer driver genes. However, there is no gold-standard method for identifying breast cancer driver genes. Therefore, this study aims to develop a model to predict high-confidence breast cancer diver genes using already-developed computational tools and already-produced breast cancer data. Primary breast cancer data were retrieved from the Cancer Genome Atlas Program (TCGA). Here we use twenty-seven different gene prediction tools that calculate each gene’s effect and variant in the primary gene dataset. The primary dataset feeds as the input for each tool. The results retrieved from each tool are recorded as the secondary dataset. The latest breast cancer driver gene set was retrieved from DriverDBv3 and included as the target attribute. Training and testing subsets were selected using k-fold cross-validation from the secondary dataset. Attributes from the secondary dataset were ranked according to their correlation with breast cancer driver genes. The ranked data were trained using different supervised machine-learning algorithms. The attributes and the learning algorithm which produced the highest classification accuracy were selected to build the new model, BReast cancer Driver (BRDriver). The new model BRDriver achieved 0.999 area under the curve and 0.999 classification accuracy on breast invasive carcinoma data. TP53 was the most predicted breast cancer drive gene (n=246) predicted by BRDriver. Interestingly, our BRDriver model predicted CDKN2A and NFE2L2 genes as new breast cancer driver genes. Further *in vivo* and *in vitro* studies are required to determine whether these genes are indeed cancer-driver genes. Many computational tools developed to identify the cancer driver genes and variants. A few methodologies were developed to combine these tools and increase efficiency for detecting breast cancer driver genes. Our BRDriver overcomes the difficulties and produces high-confidence breast cancer driver genes.

## I. INTRODUCTION

Breast cancer holds the highest number of new cancer cases (11.7%) and the fifth most cause of cancer deaths (6.9%) among the world population [1]. Breast cancer incidents and mortality drastically increased in the Asian population. It is the most diagnosed cancer (45.4%) and the most cause of cancer deaths (50.5%) in the Asian population. Genetic screening became one of the popular screening methods among breast cancer patients, which increased the monitoring and surveillance of cancer. Increased awareness of cancer screening helps with the early detection of cancer and treatment planning [2].

Driver mutations in genes cause cells to become cancer cells [3]. Distinguishing driver mutations from passenger mutations is not easy to approach [4]. Correctly identified and characterized driver genes with driver mutations can be used in cancer diagnosis [5]. New studies continued with genetic screening and produced a large amount of cancer-related genetic data. Thousands of somatic mutations were identified in these studies. Many computational tools and methods were developed using genetic data to identify the driver genes from the passenger genes [6]. Besides the classical drivers like BRCA1, BRCA2, TP53, PTEN and ATM, computational tools recently identified new driver genes such as CCND1, ERBB2, FGFR1, MYC and PIK3CA [7]. However, these single computational tools produce more biased results with less accuracy due to the heterogeneous mutation rate across the genome and false-positive calls in cancer with high mutation rates [8]. Thus, there is no gold-standard method for identifying breast cancer driver genes with high accuracy and confidence.

To that end, this study aims to build a model, BRDriver to predict high-confidence (HC) breast cancer driver genes using the existing seven different computational tools and random forest machine learning algorithm. This study retrieved data from Breast Invasive Carcinoma (TCGA, PanCancer Atlas) patient data from Cancer Genome Atlas Program (TCGA) [9]. Computational analysis is fast and less expensive compared to conventional laboratory methods. It is a more realistic option for handling high-throughput data. BRDriver will help with the early detection of breast cancer, and stakeholders can plan their treatments early.

## II. METHODOLOGY

### A. Selection of Computational tools

A literature survey was performed using specific keywords, “driver gene identification”, “driver mutation identification”, “Variant prediction” to identify the computational tools for this study. We found list of tools which predicts different features. The list includes Variant Effect Predictors, Gene Predictors, Non-Coding region effect predictors, Cancer predictors, Variant predictors and Clinical Relevance predictors. From the list, 27 computational tools were selected for this study. These tools calculate the effect of each gene and each valiant and produced a score.

### B. Preparation of Primary and Secondary dataset

Breast Cancer data were retrieved from The Cancer Genome Atlas Program (TCGA). TCGA generated data across 33 cancer types include more than 10,000 tumors. The Breast Invasive Carcinoma (TCGA, PanCancer Atlas) dataset initially selected as the Primary dataset for this study. It includes 1084 patient samples. The Primary dataset was in the MAF format and it converted into “.txt” format using a python script. The “.txt” file includes sequential columns only including Chromosome name, Chromosome Position, Strand, Reference-Base, Alternate-Base, Sample and Tag. Converted “.txt” file feeds as the data input for each selected computational tool. Results from each tool were recorded as the Secondary dataset.

### C. Retrieval Breast cancer driver gene list

The latest breast cancer driver gene set was retrieved from DriverDBv3. Incorporate with somatic mutation, RNA expression, miRNA expression, methylation, copy number variation and clinical data, DriverDBv3 included 313 breast cancer drivers from Cancer Gene Census. This gene-set included as the target attribute when training the data.

### D. Data Sampler and Ranking

Training and Testing subsets selected using k-fold cross-validation from the Secondary dataset. To include the most informative attributes from the training data, the attributes were ranked according to their correlation with breast cancer driver genes. The Information Gain score used for attributes Ranking. Ranked attributes were trained using different learning algorithms.

### E. Learning Algorithms

Five different supervised machine learning algorithms were used as the learning algorithms. K-nearest neighbours algorithm; KNN produce a value that is the average of the value of k nearest neighbours. Support Vector Machine; SVM is a classification algorithm separate classes by using a regression line. Logistic Regression is a simple and efficient classification algorithm that provides a probability score for observations. Naïve Bayes is a fast probabilistic classification algorithm based on the Bayes theorem. Random Forest algorithm gives predictions using an ensemble of decision trees.

### F. Test and Score

The results evaluated through an iterated process. The attribute ranked at first selected for the first iteration. It was trained using learning algorithms and their scores were calculated. For the second iteration, the attributes ranked at first two were selected. They were also trained using learning algorithms and their scores were calculated. The iteration process continued 27 times until all of the 27 attributes were covered. In each iteration, the Area Under Curve and the Classification Accuracy were calculated related to its learning algorithm. Finally, the ROC analysis and Confusion Matrix analysis were done for each iteration.

### G. Model Selection, Model Prediction and Model Evaluation

The iterations which produced the highest classification accuracy were identified. ROC and Confusion Matrix were observed in that iteration. The attributes and the learning algorithm from that iteration were selected for the new model building. The new model, BRDriver build using python model building module and saved as a pickle file. Five different publicly available breast cancer datasets were downloaded and they were checked using the BRDriver model. Predicted results from the BRDriver model were evaluated using the DriverDBv3.

### H. BRDriver repository

Here we present the BRDriver github repository that contained the python script and other resources. Users can clone the repository and run the program. Users need to pre-install some python packages.

## III. RESULTS AND DISCUSSION

Through the literature survey, 27 computational tools were selected for this study. Some tools predict a score and the pathogenic behaviour of the variant. Only the score was recorded.

Breast Invasive Carcinoma (TCGA, PanCancer Atlas) data, the Primary dataset consisted of 1084 patient samples. There were 84366 total mutated genes and 10662 genes with structural variants. Breast Invasive Ductal Carcinoma is the most prevalent cancer type (72%, n=780) and Invasive Breast Carcinoma (0.1%, n=1) is the least prevalent cancer type in the Primary dataset. Except for a small percentage of men (1.1%, n=12), the majority of the patients were women (98.9%, n=1072) in the Primary dataset.

The Secondary dataset attributes were ranked according to their correlation with breast cancer driver genes. Fig. 2 shows that Mutpanning produced the highest information gain score (0.126) while DANN produced the lowest information gain score (4.960×10^-5^).

**Figure 1.**
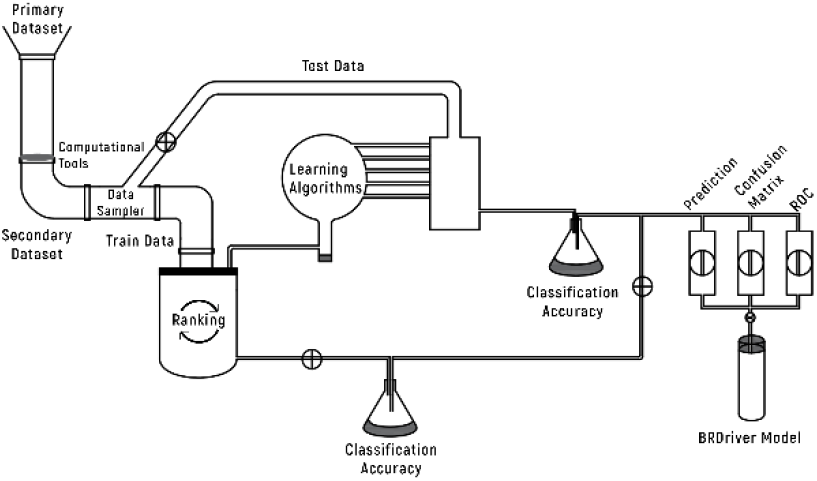
BRDriver methodology

**Figure 2.**
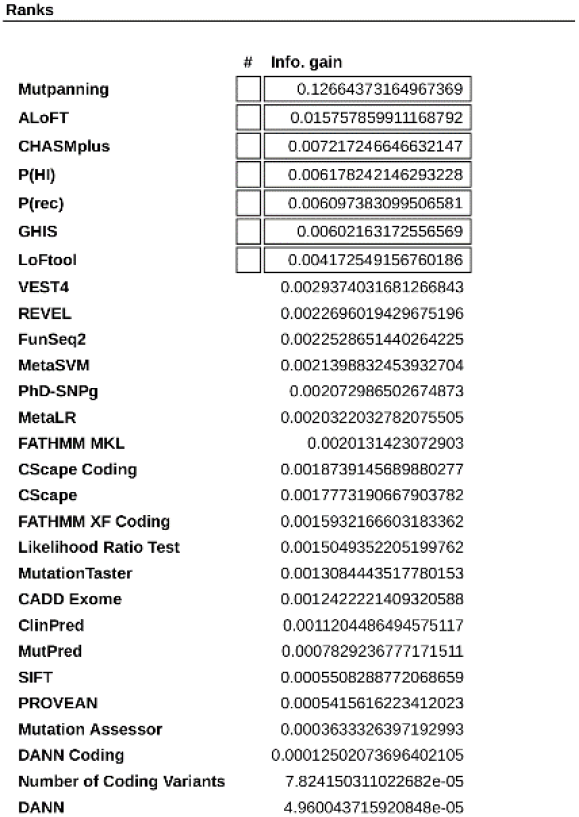
The correlation between computational tools and the breast cancer driver genes

The seventh iteration produced the highest classification accuracy (0.999) along with the highest AUC (0.999) when using the random forest learning algorithm. Fig. 3 shows the ROC curves and Fig. 4 shows the confusion matrix related to the seventh iteration and the twenty-eighth iterations. Ranked data reduced the classification errors. The seventh iteration produced the lowest miss-classified sample number (n=34).

**Figure 3.**
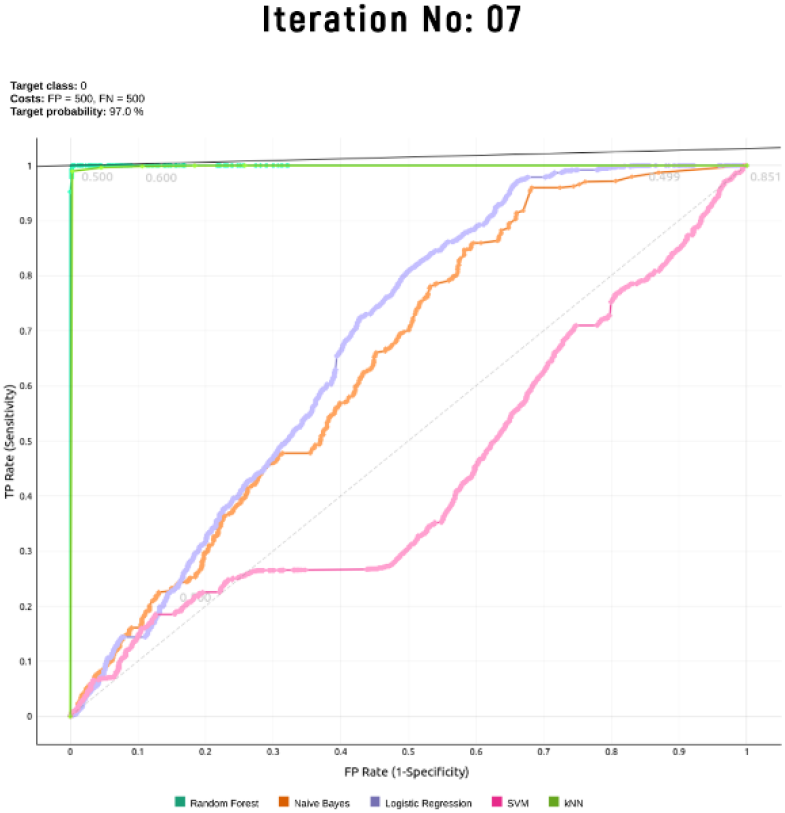
ROC curves for iteration no: 07

**Figure 4.**
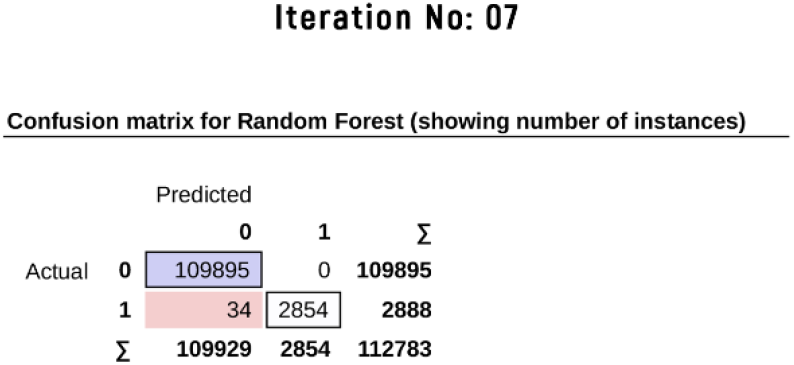
Confusion matrix for the iteration no: 07

Table 1 shows the five other breast cancer studies selected for the model prediction and model evaluation. Two novel Breast Cancer driver genes CDKN2A and NFE2L2 were identified in this study.

**Table 1.** MODEL PREDICTION

| Study | Driver Genes | Novel Driver Genes |
| --- | --- | --- |
| Breast Invasive Carcinoma (TCGA, Nature 2012) | 244 | - |
| Breast Cancer (MSK, Clinical Cancer Res 2020) | 74 | CDKN2A |
| Proteogenomic landscape of breast cancer (CPTAC, Cell 2020) | 01 | - |
| The Metastatic Breast Cancer Project (Provisional, February 2020) | 499 | NFE2L2 |
| Breast Cancer (MSKCC, NPJ Breast Cancer 2019) | 137 | NFE2L2 |

Cancer driver gene identification is much needed for accurate oncological analysis. Many computational tools were developed to identify the cancer driver genes and variants but very few methodologies were developed to combine these tools and increase their prediction accuracy. This study uses two main types of computational tools, Gene predictors and Variant predictors. A few gene predictors are available for a feasible study. GHIS, LoFtool, P (rec), and P (HI) tools are the gene prediction tools used in this study. The GHIS database provides the haploinsufficiency score for a given particular gene. It integrates with a large dataset and predicts results. [10]. Based on the loss of function, intolerance and tissue expression of a given gene, LoFtool provides a score for a given gene. [11]. P(rec) estimate the probability of a given gene is a recessive disease gene or not [12]. P(HI) database provides Haploinsufficiency score for a given gene [13]. The rest of the tools are variant predictors. Other than the tools, the number of coding variants calculated for a given gene includes in the Secondary dataset.

When combining different tools for a particular study it is really difficult when they need different input data types and when they produce different result types. The tools selected for this study use same format of input data type. They needed Genome annotation, Chromosome Position, Strand, Reference-Base, Alternate-Base for giving their predictions.

This study proposed the BRDriver model for the identification of breast cancer driver genes. BRDriver built upon seven different variant prediction tools Mutpanning, ALoFT, CHASMplus, P (rec), P (HI), GHIS, LoFtool and the Random Forest machine learning algorithm.

### A. Novel breast cancer driver genes

In this study, BRDriver identified two new potential breast cancer driver genes CDKN2A and NFE2L2. Table 1 shows the 05 selected independent studies with BRDriver predictions.

#### CYCLIN-DEPENDENT KINASE INHIBITOR 2A; CDKN2A

CDKN2A act as tumor suppressor gene. It encodes the p16^INK4a^ protein. This protein plays a major role in cell cycle. Mutated CDKN2A gene will altered the cell cycle regulation. [14].

#### NUCLEAR FACTOR, ERYTHROID 2 LIKE 2; NFE2L2

NFE2L2 act as a transcriptional factor which regulates oxidative stress and inflammatory response. A positive correlation was found between NFE2L2 and breast invasive carcinoma. [15]

## IV. CONCLUSION

This study proposed the BRDriver model for breast cancer driver gene identification. BRDriver built upon seven different variant prediction tools and it uses the Random Forest machine learning algorithm for giving predictions. BRDriver needs the chromosome number, chromosomal position, Reference Allele and the Alternative Allele information for a particular gene.

